# Selective phosphorylation of AKT isoforms in response to dietary cues

**DOI:** 10.1101/693242

**Authors:** Laura Christin Trautenberg, Elodie Prince, Cornelia Maas, Nora Beier, Freya Honold, Michal Grzybek, Marko Brankatschk

## Abstract

A calorie-rich diet is one reason for the continuous spread of metabolic syndromes in western societies. Smart food design is one powerful tool to prevent metabolic stress, and the search for suitable bioactive additives is a continuous task. The nutrient-sensing insulin signaling pathway is an evolutionary conserved mechanism that plays an important role in metabolism, growth and development. Recently, lipid cues capable to stimulate insulin signaling were identified. However, the mechanistic base of their activity remains obscure to date. Here, we show that specific AKT/ Protein kinase B isoforms are responsive to dietary lipid extract compositions and potentiate cellular insulin signaling levels. Our data add a new dimension to existing models and position *Drosophila* as a powerful model to study the relation between dietary lipid cues and the insulin induced cellular signal cascade.

## Introduction

The insulin signaling (IS) pathway is evolutionary highly conserved. Alike vertebrates, *Drosophila* expresses one insulin receptor (dInR) and the downstream signal cascade shows high molecular and functional conservation (1). The dInR is a dimeric type-I membrane protein and its intracellular domain resembles the structure of a tyr-kinase (2). Bound to insulin, the dInR recruits adapter proteins and activates the PI-3 kinase (3). The PI-3 kinase converts the inner leaflet membrane lipid PI(4,5)P_2_ into PI(3,4,5)P_3_; and PI(3,4,5)P_3_ is necessary to enrich the Protein kinase B (PKB also known as AKT) (4).

Different AKT isoforms have been identified. Mice or fruit flies express three different protein versions (5,6). In *Drosophila*, the presence of individual AKT isoforms (dAKTs) is stage-dependent. Adult flies express two dAKT proteins that are different in size: the smaller dAKT^66^ and the larger dAKT^85^, close to 66kDa and 85kDa, respectively (5). Alike vertebrates, both dAKTs have two potential phosphorylation sites, dAKT^Ser505^ and dAKT^Thr342^. The functional relevance of the vertebrate AKT1^Ser473^ (which corresponds to the dAKT^Ser505^) in IS is well studied (7). The biological role of the vertebrate phosphorylated AKT1^Thr308^ (which corresponds to the dAKT^Thr342^) is more obscure, but it is widely accepted that it promotes the activity of the enzyme (7,8). dAKT is a negative regulator of the transcription factor dFOXO. When IS levels are low, dFOXO translocates from the cell cytoplasm into the nucleus (9). Whereas high dAKT activity prevents dFOXO to enter the nucleus, and the cells predominantly perform catabolic metabolism by building stocks of storage molecules such as fatty acids (10).

*Drosophila* feed preferentially on rotting fruits, a diet composed by plant material and microbes such as yeast. Fruits are the main source for carbohydrates while microbes provide dietary amino acids. It was shown that a calorie-rich diet supplemented with yeast lipids increases circulating *Drosophila* insulin like peptides (dILPs) and facilitates high systemic IS levels (11). High IS stimulates the proliferation and developmental rate of flies. Specific lipids are known to change IS in vertebrates, but their impact on insulin-secreting cells remains unclear (12,13). However, absorbed dietary lipids could change the fly lipidome as well as the presence of the dInR at the plasma membrane, or modulate the amount of PI(3,4,5)P_3_ required to recruit and stabilize dAKT.

Here, we report the isolation of *Cystobasidium oligophagum*, an ubiquitous *Basidiomycota*, from *Drosophila* droppings. Although *C. oligophagum* attracts adult flies, we show that these yeasts do not promote fruit fly fecundity or development. Compared to baker’s yeast (*Saccharomyces cerevisiae*), *C. oligophagum* produce similar amounts of protein and sugar but differ in their lipid composition. Calorie-rich food based on either yeast type supports the generative cycle of *Drosophila*, however, only diet manufactured from stationary *S. cerevisiae* accelerates its developmental rate and increases egg production. Moreover, we demonstrate that flies kept on stationary *S. cerevisiae* food upregulate selectively the dAKT^85^-isoform, and that the enzyme is highly phosphorylated. Thus, we speculate that lipid cues derived from stationary *S. cerevisiae* stimulate the accumulation of PI(3,4,5)P_3_ at the inner plasma-membrane leaflet of *Drosophila* cells. Due to its PH domain especially dAKT^85^ binds to PI(3,4,5)P_3_ and locally trapped, is activated by the phosphoinositide-dependent protein kinase (PDK1) and the rictor-mammalian target of rapamycin complex 2 (mTorC2) (14,15).

## Results

### *Cystobasidium oligophagum* and *Saccharomyces cerevisiae* have different lipid qualities

On rotting plant material, yeast represent the dominant microbes that are preferred by *Drosophila*. To investigate what fungi are associated with flies in the laboratory, microbial isolates from fly droppings were analysed. Fly poo positioned on plant food at 20°C contained *C. oligophagum*, but not *S. cerevisiae*. To compare the caloric value of either yeasts, *C. oligophagum* or *S. cerevisiae* were cultivated at 20°C in a defined medium. The fungi were harvested in their exponential (EP) and stationary growth phases (SP) (Fig.S1D, E), and subsequently, their protein and trehalose content was quantified by using commercial detection assays. Remarkably, both fungi produce very similar amounts of protein and trehalose (Fig.S1F, G). To evaluate fungal lipids, yeast lipids have been extracted and total lipid amounts were normalized. Individual lipid classes were separated by using reverse phase Thin-Layer Chromatography (2D-TLC). The growth state of *S. cerevisiae* has shown to be important for the quality of at least one polar lipid class (Fig.S1Ha, b). Moreover, *S. cerevisiae* that reached the SP produce additional neutral lipid classes (Fig.S1Hc,d). In contrast, the lipid profiles from *C. oligophagum* show minimal growth-dependent quality changes (Fig.S1I). Although lipid qualities are well assessable using 2D-TLC, relative quantities are hard to estimate. To value relative amounts of neutral lipids, lipid profiles were analyzed using 1D-TLC. Standardized to phosphate, the lipidome of *C. oligophagum* is overrepresented by lipids with properties shown by free fatty acids. Moreover, the relative amounts of individual lipid classes do not vary between EP and SP. In contrast, EP *S. cerevisiae* have less triacylglycerids (TAGs) and sterol-esters in comparison to SP yeast. In addition, *S. cerevisiae* have less neutral lipids compared to *C. oligophagum* (Fig.S2A).

To show the lipid distribution in fungal cells, live yeast were stained with Bodipy-505. The dye is capable to penetrate the plasma membrane and it preferentially accumulates in lipid-rich regions such as lipid droplets. As expected, in *S. cerevisiae* cells, Bodipy-505 is enriched in lipid droplets and it was noted that SP yeast cells contain a higher number of such organelles (Fig. 1A, B). However, the lipid distribution of *C. oligophagum* cells is rather static and does not change between EP and SP (Fig.1C,D). Taken together, *S. cerevisiae* lipid extract qualities are dependent on their growth phase; whereas *C. oligophagum* lipid profiles appear constant.

**Figure 1:**
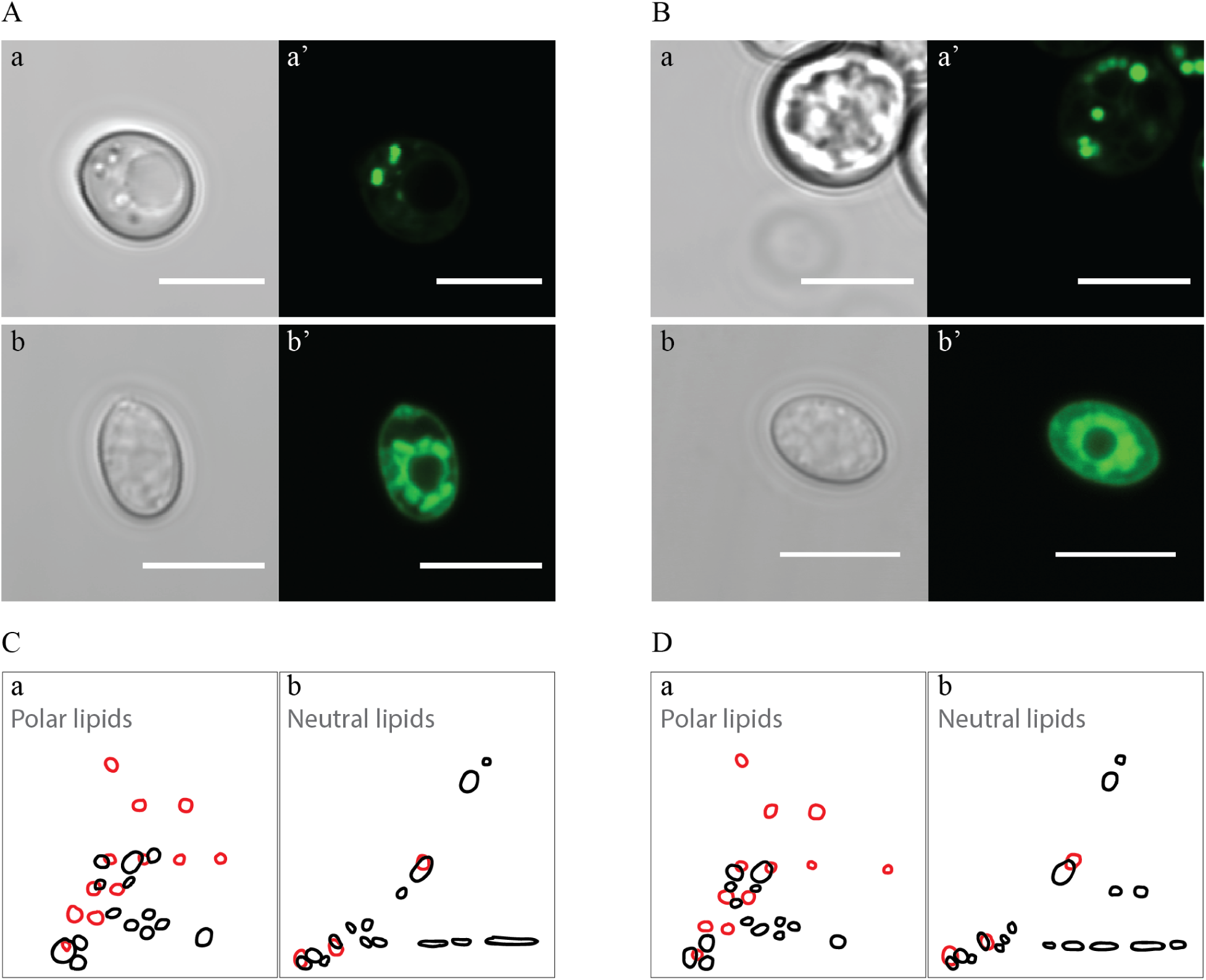
*Cystobasidium oligophagum* and *Saccharomyces cerevisiae* have different lipid profiles. A,B Photographs of yeast harvested at specific growth stages probed with BODYPI-505. Shown are pictures taken with DIC (Aa,b and Ba,b) or fluorescence (Aa’,b’ and Ba’,b’) from *S*.*cerevisiae* in exponential (Aa,a’)and stationary growth phase(Ba,a’) and *C. oligophagum* in exponential (Ab,b’)and stationary growth phase(Bb,b’). Scale bar indicates 5µm. C,D Shown are 2D-TLC profiles of polar (Ca, Da) and neutral lipids (Cb,Db). Lipid signatures from *S*.*cerevisiae* (black) and *C*.*oligophagum* (red) from exponential (C) and stationary (D) growth stages are overlayed.

### Designed yeast food mimics the biological activity of the respective yeast

To test if these fungi attract adult flies, wild-type *OregonR* were placed in feeding chambers to video-record the fly behaviour. On opposing sides of the chamber, two plant food baits were put as a control (Fig. 2A). This experimental setup did not reveal any biased feeding. Next, the experiment was repeated however, this time one plant food bait was exchanged with yeast. Both yeast, *S. cerevisiae* and *C. oligophagum*, strongly attracted adult flies after a short adaptation time (Fig. S1A-C). Females deposed their eggs preferentially close to the yeast (Fig.2B, C). Regarding the egg laying rate, it was noted that flies feeding on *S. cerevisiae* produced reproducibly higher egg numbers compared to flies feeding on *C. oligophagum* or plant material only (n=5, total egg number: *C. oligophagum*=299, *S. cerevisiae*=578 and plant material=184 eggs, each assay plate with 20-30 females and 15 males). The fecundity of females is a direct readout for their metabolic activity. Thus, the different reproductive activity could be explained by a very different caloric content or nutritional profile of both provided yeast species.

**Figure 2:**
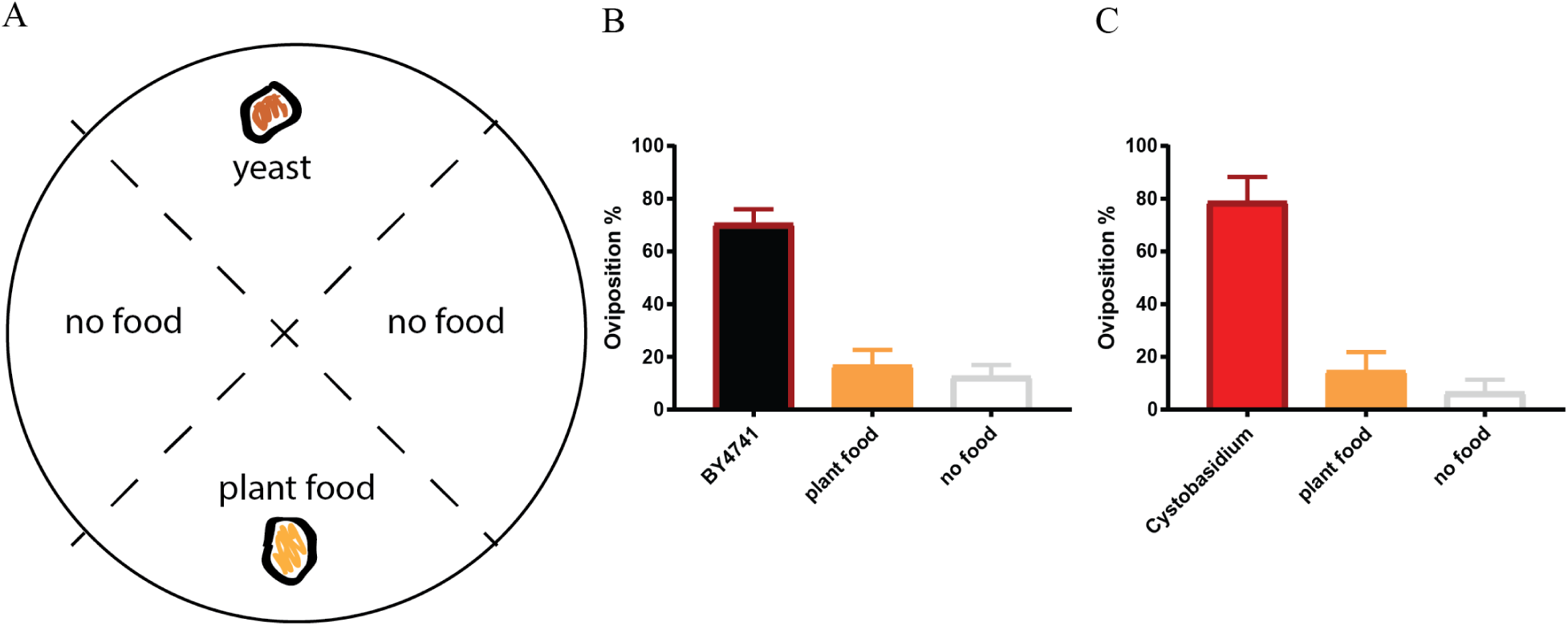
*Cystobasidium oligophagum* attracts to *Drosophila melanogaster*. A Scheme that depicts our quantification approach, assay plates are divided into four sectors loaded with food (yeast or plant food) or without a bait (no food). B,C Plotted are percentages of eggs positioned in different food sectors: *S. cerevisiae* (B in black), *C. oligophagum* (C in red), plant food (B,C in yellow) or no food (B,C in grey).

To neglect possible caloric differences between *C. oligophagum* and *S. cerevisiae*, four different equi-caloric food recipes (approx. 550 kcal/l) were designed. All diets are rich in proteins and sugars resulting in approximately identical carbohydrate: protein (2:1) ratios, and therefore preventing carbon or amino acid shortages (Tab.1). Due to the nature of our food preparation protocol, yeast proteins or sugars form only a small proportion, while the fungal lipidomes are the only lipid source of our produced food types. To estimate the quantity of lipids in the food, the food’s dry mass was weighted, and phosphate was measured from respective lipid extracts. Food (F) produced from *C. oligophagum* (*C*.*o*.) or *S. cerevisiae* (*S*.*c*.) show similar lipid mass (n=3, *C*.*o*.-EP-F=0.6; *C*.*o*.-SP-F=0.5, *S*.*c*.-EP-F=0.3 and *S*.*c*.-SP-F=0.4 mg phospholipids/g food). Therefore, the lipids from both yeasts contribute with almost similar caloric amounts (even if total lipids divert 10 times from the measured amount of phospholipids the caloric difference in the food recipe would be lower than 20cal/l). However, the heat treatment during food preparation could break some yeast products. To analyse the lipid profiles of the designed diets, the food lipids were extracted, and the phosphate-standardized samples were analysed. Interestingly, respective food samples showed profound differences if compared to yeasts lipid extracts themselves. TAGs and sterol-esters are reduced in the designed diets, while sterols are enriched (Fig.S2B, C). Taken together, the lipid profiles of the designed diets differ from respective yeast lipidomes.

To test if the created yeast food still induces the physiological changes of yeast feeding *Drosophila*, the feeding-behaviour experiments were repeated. Given the choice between plant material and yeast-based diets, adult flies tended to feed on the latter. Food produced from *S. cerevisiae* grown to SP elevated the fecundity of adult females enormously (n=9; total egg number: *C*.*o*.-EP-F=135, *C*.*o*.-SP-F=150, *S*.*c*.-EP-F=184, *S*.*c*.-SP-F=291 and plant material only=72 eggs, each assay plate 9 females and 3 males). Thus, the designed yeast diets mirrored the activity of the respective living yeast. To investigate if food-associated microbes or microbial metabolites modulate larval growth and survival, microbe-free (axenic) animals were reared. Axenic animals rely entirely on provided food compositions and show a direct response to nutritional settings. Interestingly, only larvae kept on *S*.*c*.-SP-F were able to match the developmental speed and success of their microbe-bearing siblings. Moreover, all other microbe-free cultures developed slower and with lower survival rates than larvae fed with *S*.*c*.-SP-F (Fig. 3).

**Figure 3:**
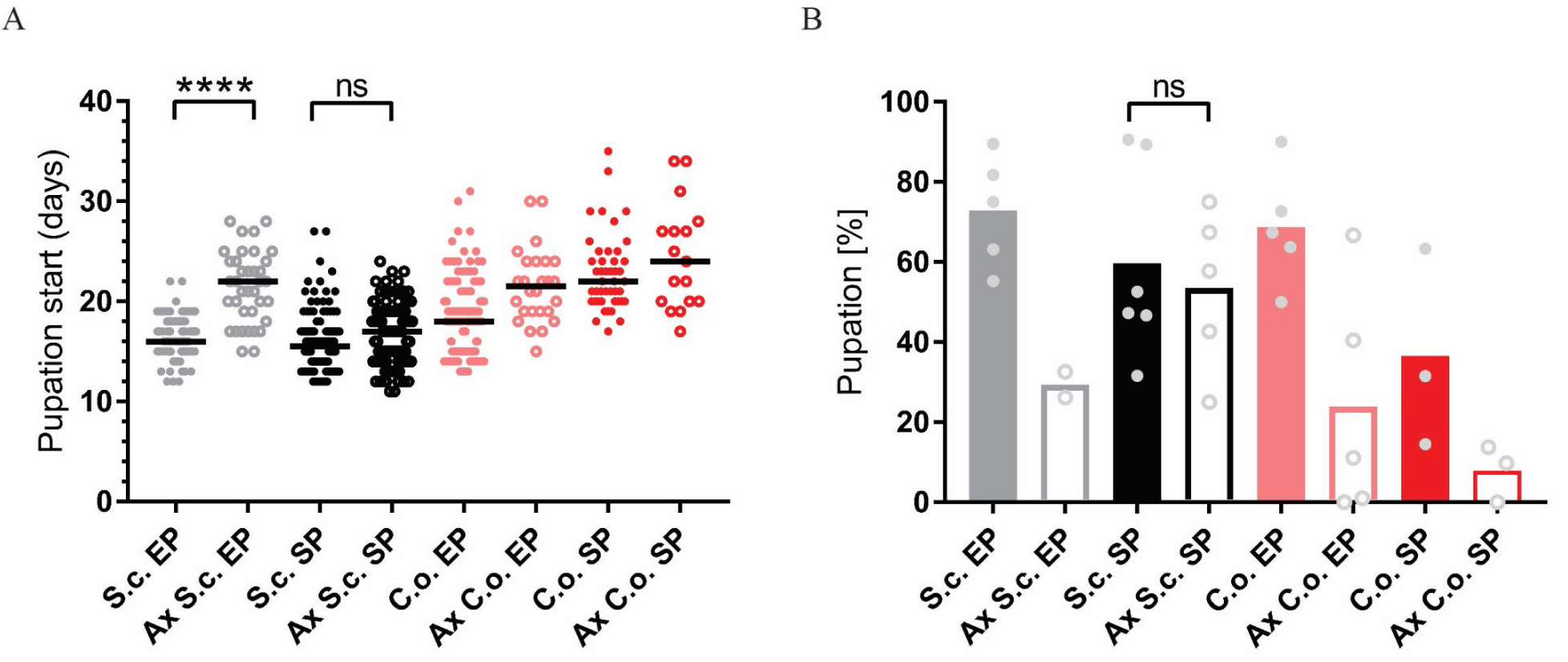
Stationary yeast food rescues the metabolic syndrome of axenic larvae. A,B Plotted is the developmental speed (A) and survival rate (B) of axenic (AX) and microbe-associated larvae kept on food based on exponential (EP) and stationary (SP) *S*.*cerevisiae* (S.c.) or *C*.*oligophagum* (C.o.). Each spot represents one tracked individual (A) or one experimental cohort of minimum 20 individuals (B). Significance was calculated using ANOVA, n.s. equals none significant, **** equals p<0.0001.

In sum, the designed food types are indifferent in their caloric content, their carbohydrate: protein ratios are identical, and the diets are rich in amino acids and monosaccharides. In consequence, the quality and quantity of the respective dietary lipid load is modulating the response of flies.

### Dietary lipid extracts regulate the cellular insulin signal cascade

Animals fed with high-fat diet or kept on chow-food show very different lipidomes (16). We have shown that the lipid extract composition of the different yeast foods varies, and we wondered if the nutritional lipid quality in the experimental setups would induce endogenous lipid changes in feeding *Drosophila*. To do so, adults were transferred from normal food onto experimental foods for 14 days. Lipid extracts from larvae or adult fly heads did not reveal changes in the composition of complex endogenous lipids (Fig.S3). At this point, it is not possible to exclude the possibility that structural qualities of absorbed and integrated dietary fatty acids regulate the fecundity and developmental rate. However, it is shown that different lipid profiles do not change cellular membrane properties at set experimental temperatures (17). On the other hand, *S. cerevisiae* are able to facilitate systemic IS in fruit flies (11). To test if larvae, kept on the designed diets, modulate their IS cascade, the localization of the transcription factor dFOXO was analysed. dFOXO is the most-downstream target of the insulin pathway and resides in the nucleus at low metabolic rates. However, rich nutrition typically capable to foster IS deactivates dFOXO, and the protein translocates into the cytoplasm (9). Like expected, only fat body cells from larvae kept on *S*.*c*.-F show a predominant cytoplasmic dFOXO localisation (Fig. 4A). Many different dILPs regulate the larval development and thus, mimic the function of vertebrate insulin-like growth factors (IGFs) (1). Moreover, invertebrate dILPs probably represent an ancestral regulative network that channels IGFs and insulin-like signalling to one receptor (28,29). In consequence, developmental aspects blur the larval metabolic insulin signalling. To circumvent the problem, we decided to analyse the metabolic cellular insulin cascade in adult flies.

**Figure 4:**
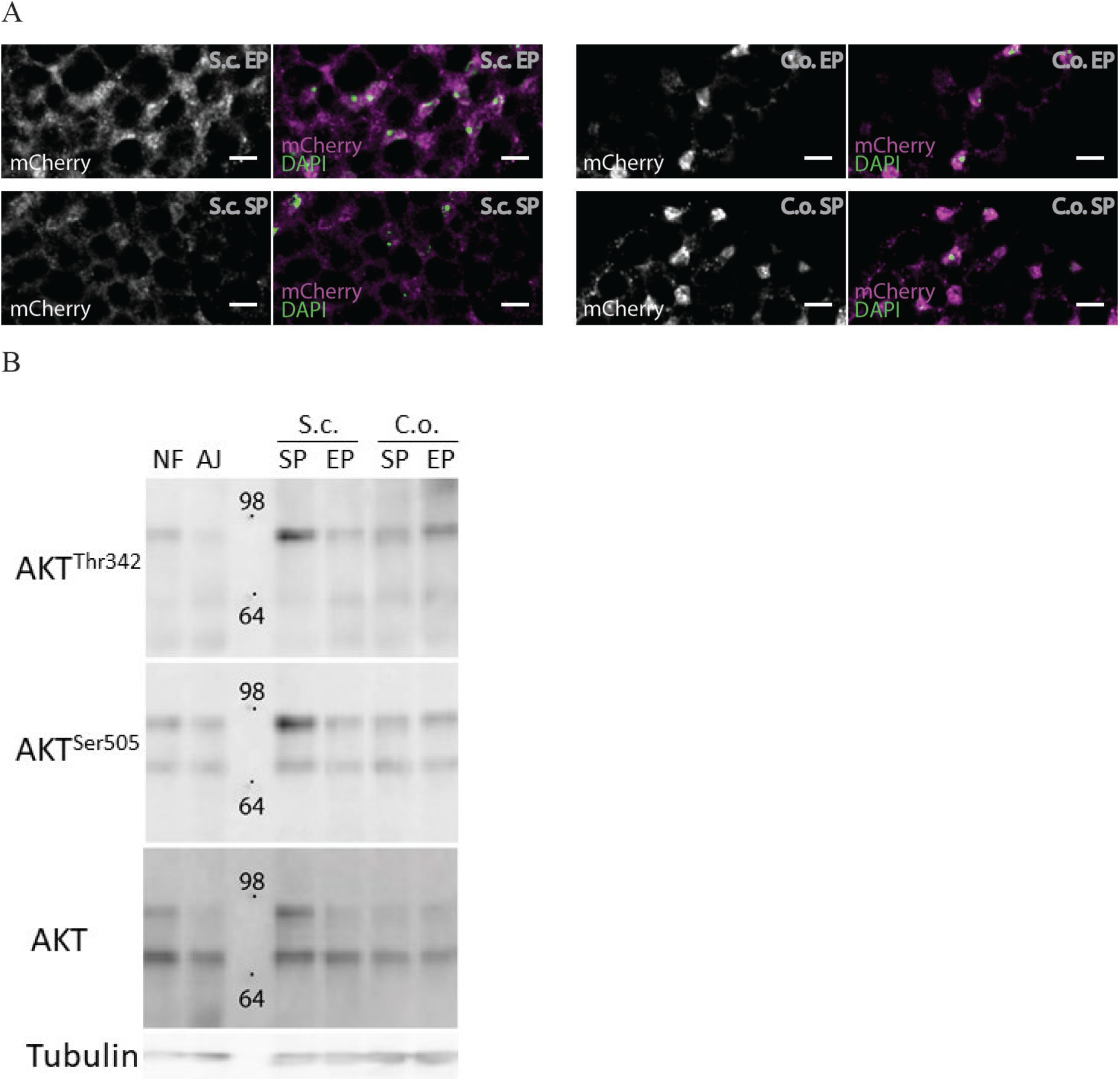
Individual dAKT isoforms facilitate dInR induced cellular insulin signaling. A Shown are photographs from larval fat body cells stained with DAPI (green) and probed for dFoxo-mCherry (white/ magenta) from animals kept on food produced from *S*.*cerevisiae* (S.c) or *Cystobasidium oligophagum* (C.o.) harvested in the exponential (EP) or stationary growth phase (SP). Scale bars indicate 10µm. B Photographs of a Western-blot membrane with protein lysates from adult heads of flies kept on normal food (NF), apple juice agar (AJ), food from stationary (SP) and exponentially (EP) grown *S*.*cerevisiae* (S.c.) or *C*.*oligophagum* (C.o.). The membrane was probed for phosphorylated dAKT (dAKT^Thr342^ and dAKT^Ser505^), total expressed dAKT protein (AKT) and Tubulin (Tub). Depicted is the position and size (in kDa) of marker proteins.

The upstream regulator of dFOXO is the Protein kinase B/dAKT. Phosphorylated active dAKT deactivates dFOXO. Therefore, we speculated that only flies kept on *S*.*c*-F should show high dAKT phosphorylation levels. Adult flies were bred on normal food and then transferred onto the different yeast-based diets for 14 days at 22-23°C. The two reported isoforms of dAKT were detected (Fig. 4B) (5): relative high amounts of the lower mass dAKT^66^ and traces of the larger dAKT^85^ protein. Astonishingly, only in samples from flies kept on *S*.*c*.-SP-F dAKT^85^ levels matched almost the presence of dAKT^66^ protein. Moreover, we found that only dAKT^85^ is phosphorylated at both positions (*i*.*e*. Ser^505^ and Thr^342^) on all diets. Thus, we conclude that in adult differentiated cells, dAKT^85^ is the predominant active AKT isoform and that dietary lipids present in *S*.*c*.-SP-F diet elevate dAKT^85^ amounts to facilitate cellular IS.

## Discussion

Environmental factors drive the spread and the dynamic behaviour of insect populations as well as the quality and abundance of accessible food sources. Recent findings add another dimension to this simple equation, the identity and state of food-associated microbes. *Drosophila* are attracted by volatile yeast cues and prefer to feed on calorie-rich rotting plant material infested by yeast and other microbes. Such microbes share often the optimal temperature range of fruit flies (18–20). Hence, available rotting fruits or other decomposing plant materials in a given area are infested by different yeast species. Indeed, flies caught in the wild were associated with different yeast, however, the microbial environment was strongly dependent on local environmental factors such as food source or temperature (21).

It was shown that lipid products from *Saccharomyces cerevisiae* accelerate the generative cycle of flies by increasing the systemic insulin signalling rate (11). However, *S. cerevisiae* are not predominant in natural habitats and it is likely that the interaction between fruit flies and their microbial surrounding is more complex (22). In consequence, several questions arise in our quest to understand the physiology and evolutionary conserved metabolic circuits of fruit flies.

At first, we attempted to answer the question if all food-associated yeast provide factors that support *Drosophila*. To do so, we isolated and identified *Cystobasidium oligophagum* from fly droppings. *C. oligophagum* grows on decaying plant material and are ubiquitous (23). Our behavioural assays show that alive *C. oligophagum* attract adult flies and that flies ingest the yeast. Thus, *C. oligophagum* is a food-associated fungus that interacts with wild fruit flies however, it is unlikely a stable and lasting microbe-host interaction.

Flies kept on plant material supplemented with *C. oligophagum* show a much lower fecundity compared to flies feeding on *S. cerevisiae*. Our findings allow for different explanations, either *C. oligophagum* is a poor diet or *C. oligophagum* products do not stimulate the metabolism of *Drosophila*.

We have compared the protein, sugar and lipid content of *C. oligophagum* with yields present in *S. cerevisiae*. Total protein and sugar levels are very similar for both fungi, whereas *C. oligophagum* is richer in fatty acids (23). We concluded that the caloric fungal value is unlikely responsible for the low biological activity of flies kept on *Cystobasidium*.

Since *S. cerevisiae* lipid extracts modulate the IS in *Drosophila* (11), we decided to focus on *C. oligophagum* lipids and neglect the value of fungal proteins or sugars by creating different diets. We show that the lipidomes of our diets differ from the lipid profiles of the corresponding yeast. Nevertheless, repeated behaviour and egg laying assays consolidated our initial findings with the living fungi. Although rich in calories, *C. oligophagum* food did not accelerate the proliferation rate of fruit flies with respect to *S. cereviciae*. However, examining the metabolic activity of dietary lipids is not as trivial as it might seem at first glance.

Fly-associated microbes can modulate the nutritional input and, in some cases, promote directly or indirectly systemic insulin signalling. For instance, *Acetobacter pomorum* secrete metabolites capable to induce insulin signalling (24). In poor nutritional conditions, *Lactobacterium* replenish amino acid shortages and thus, restore the metabolic activity of the host (25). To exclude the microbial variable in our experiments we decided to use heat-sterilized diets supplemented with fungal resistance factors.

Axenic (microbe-free) animals tend to develop a metabolic syndrome similar to insulin resistance resulting in lower survival and slower developmental rates (24,26). Therefore, we decided to run microbe-bearing and microbe-free cultures in parallel. Only axenic cultures thriving on stationary *S. cerevisiae* food (*S*.*c*.-SP-F) matched the developmental parameters from siblings kept on respective microbe-infested diet. On the other hand, living microbes in general rescued the hampered *Drosophila* development on all tested food types.

This finding provokes the idea that lipid extracts from stationary *Saccharomyces* might induce the same metabolic response as many living microbes. At this point, we miss solid evidence to address the issue in deep. However, the observation that microbe-associated and axenic animals kept on *S*.*c*.-SP-F are indifferent argues for one identical rescue mechanism.

Our cultivation protocol stimulates the larval digestive system mildly and food-associated microbes likely induce an immune response. It is widely accepted that the Toll-like signalling, one integral component of the *Drosophila* immune system, is activating the kinase AKT (27). dAKT in turn regulates dFOXO, the downstream target of the insulin signalling cascade and the cellular uptake of sugars and amino acids (9). Taken together, the activation of the Toll-like pathway may phenocopy the function of the activated dInR.

In *Drosophila*, yeast lipids are able to increase the production of dILPs by insulin producing cells (11) and the genetic increase of circulating dILPs is capable to increase systemic IS. Thus, it is likely that the immunological response to microbes and yeast lipids modulate the insulin cascade at different levels to the same result.

Vertebrates express different AKT isoforms, for instance mice or humans express three different AKT proteins. Surprisingly, knock out mice with only one functional copy of AKT1 are viable but show some physiological changes and are sensitive to dietary sugar loads (6). However, most studies presume a parallel activity of all AKT isoforms and value the contribution of individual AKT isoforms in the insulin signalling cascade based on their present levels or compartmentalization (6,30). Although evidence in flies is missing, the spatial organization of different dAKT isoforms could account for the detected differences in phosphorylation and protein levels, yet it remains elusive how cells could realize such a transient regulation.

Alike vertebrates, fruit flies express three different isoforms of the kinase AKT (5). Like reported, we have detected two isoforms in our samples, dAKT^66^ and dAKT^85^. Regardless of the provided food, both dAKT isoforms show a solid phosphorylation at the position dAKT^Ser505^. However, only dAKT^85^ is in parallel phosphorylated at the position dAKT^Thr342^. In addition, we show that protein levels of dAKT^85^ are elevated in flies fed with food from stationary *S. cerevisiae*. Therefore, our working model predicts that lipid extracts from stationary *S. cerevisiae* increase circulating dILP levels and more insulin peptides bind to their receptor (11). The insulin bound dInR recruit additional PI3 kinases, which induce the production of Phosphoinositol-(3,4,5)-phosphate (PIP_3_) by conversion of membrane integrated Phosphoinositol-(4,5)-phosphate (PIP_2_) (4). Therefore, the PIP2:PIP3 ratio is shifted and more cytoplasmic dAKT^85^ associates with PIP3 and is phosphorylated by local kinases. Further, phosphorylated dAKT^85^ regulates the downstream transcription factor dFOXO, which would implicate that dAKT^85^ is responsible for the insulin mediated glucose disposal. Interestingly, our western blot data indicate that the major pool of the dAKT^66^ protein is inactive. One reason for our finding could be that dAKT^66^ is confined and not accessible in differentiated cells. We reinforce the idea that dAKT^66^, like in vertebrates AKT3, fulfils more developmental roles (5,30) and is not engaged in the metabolic response of adult flies (Fig. 5).

**Figure 5:**
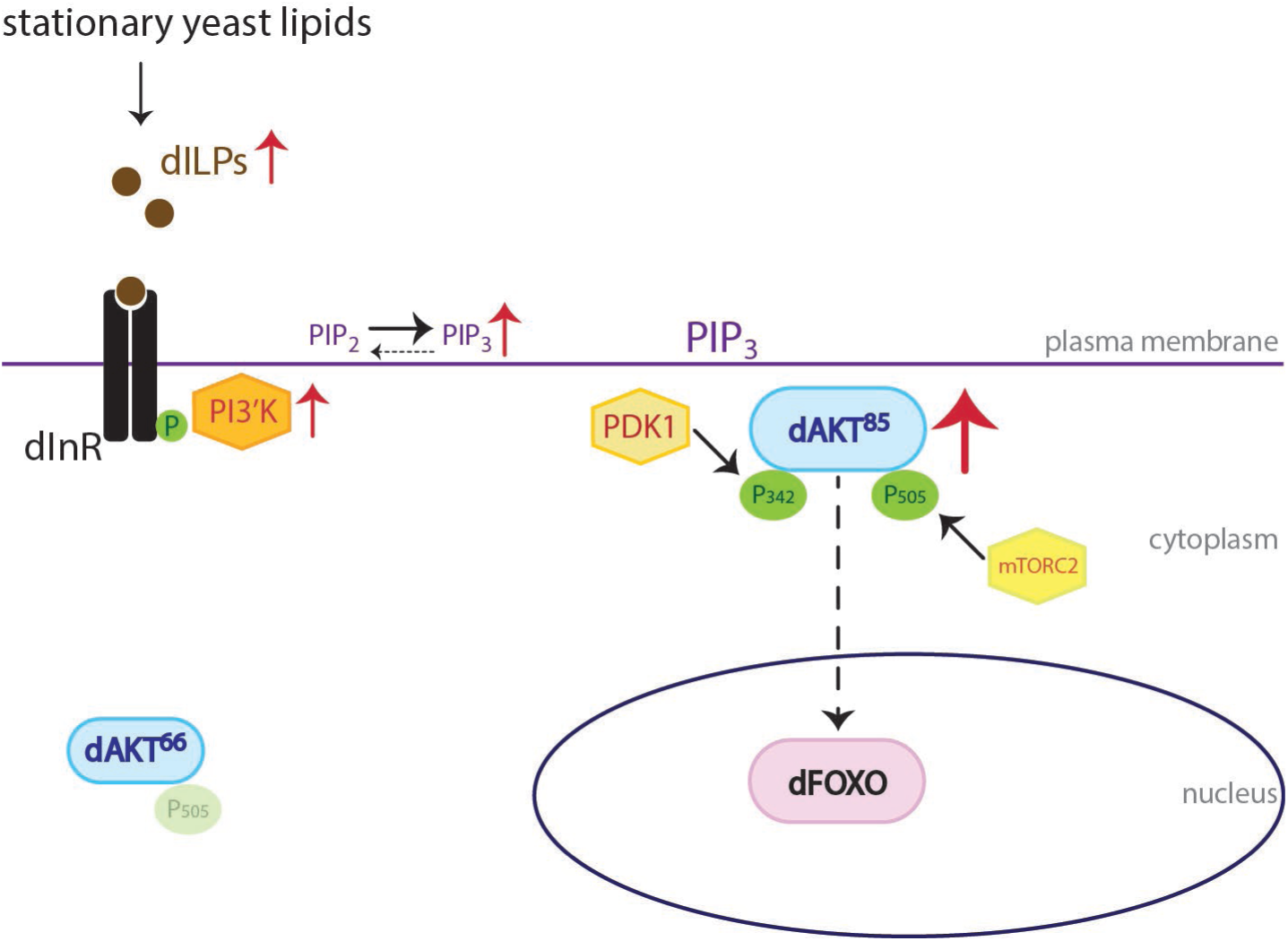
Transient amplification of the cellular insulin response. Model that depicts the transient activation of different dAKT isoforms in response to the activity of the insulin receptor.

In sum, we add to the complexity of microbe-host interactions the variable microbial growth stage, and we spotlight the differential phosphorylation and expression of dAKT isoforms as a versatile tool to facilitate the cellular insulin signal cascade.

## Acknowledgements

We are thankful to Martin Kaltenpoth (University Mainz) for identifying *Cystobasidium oligophagum*, experimental work by LT, EP, CM, FH, NB, MG and manuscript concept by LT, EP, MG and MB. This work was supported by the German research council (DFG) by grants to MB, EP, FH, NB and LT (BR5492, FOG2682-BR5493).

## Material/Methods

### Fly stocks

If not stated otherwise flies were kept at RT in a day/night cycle. *mCherry::FOXO* from S. Eaton, and *oregonR* were purchased from Bloomington stock center.

### Yeast

*Saccharomyces cerevisiae* (BY4741) was obtained from K. Ostermann and *Cystobasidium oligophagum* are wild isolates identified by M. Kaltenpoth.

### Food recipes

Normal food (https://bdsc.indiana.edu/information/recipes/bloomfood.html) and plant food (16) were produced following published protocols using dry-yeast purchased from the grocery discounter Kaufland. Saccharomyces (BY4741) and *C. oligophagum* foods were produced based on pelleted yeast obtained from SCP-medium (1.9 g/L yeast nitrogen base, 5 g/L ammonium sulfate, 20 g/L glucose, 20 g/L peptone) cultures grown at 20°C (v=100ml; wet yeast-pellet=9,04g, glucose=6g, soy peptone=2g, sucrose=3g, agar-agar=1g and nipagin=0.4g). The Exponential Growth Phase (EP) is defined at the optical density OD_600 EP_ =2-3 for *S. cerevisiae* or a mass^EP^ of 0.02g/ml for *C. oligophagum* and the Stationary Growth Phase (SP) at OD_600_^SP^>5 (*S. cerevisiae*) and a mass^SP of^ >0.04g/ml (*C. oligophagum*). Of note, *C. oligophagum* cells tend to aggregate preventing a conventional optical density measurement.

#### *Cystobasidium oligophagum* isolation

*OregonR* were kept in an open cage on plant food without resistance factors for 2 weeks at 20°C, and subsequently transferred onto apple-juice plates. Deposited droppings were resolved in water and plated onto YPD-culture media plates. After 72h at 20°C microbial colonies were picked, cultivated in YPD media and samples send for 18S ribosomal sequencing.

### Behavior assays

Flies were raised on normal food, adults transferred for 6h on apple juice plates at 22°C, and subsequently placed on assay plates (20% apple juice, 1% agar) fitted with food baits opposing each other. Fly behavior was recorded for 3h at 22°C and afterwards plates were kept for 24h at 20°C. Subsequently, flies were removed, and deposed eggs counted.

### Larval developmental tracking

Eggs collected from apple juice agar plates were washed, bleached and transferred onto apple juice plates. Later, hatched first instar larvae were placed onto food and kept at 20°C.

### Axenic animals

Eggs collected from apple juice agar plates were washed, bleached and transferred onto sterile apple juice plates. Later, hatched first instar larvae were placed under sterile conditions onto microbe-free food using aseptic tools and kept at 20°C.

### Trehalose measurements

Samples were autoclaved and processed like recommended by manufacturer (Trehalose Kit, Megazyme).

### Protein estimation

Samples were autoclaved and processed like recommended by manufacturer (BCA Kit, Pierce).

### Lipid extraction and TLC

Tissue samples were thawed on ice, homogenized in HBS using an IKA ULTRA-TURRAX disperser (level 5, 1min), and lipid-extracted by the BUME method (31). Extracted lipids were stored in chloroform / methanol (2:1) solution at −80°C.

### Biochemistry

Adult fly heads: flies were raised on normal food and adults transferred for 14 days on respective diets at 22°C (food renewal in regular intervals). Adults were snap-frozen with liquid nitrogen and fly heads removed for biochemistry. Antibodies used to probe by western blotting are AKT-Ser505 (Cell Signaling), AKT-Thr308 (Invitrogen), AKT (Invitrogen), mCherry (Cell Signaling) and Tubulin (Thermo Fisher). Larval extracts: First instar larvae were transferred on respective food and kept at 20°C. For immunohistochemistry, larvae were fixed in 4% PFA and stained with DAPI. *mCherry*-FOXO signal in larval fat body was detected using antibodies against mCherry and Alexa 555-conjugated secondary antibody (Thermo Fisher) (Zeiss LSM700). Yeast were stained with Bodipy 505/515 (Thermo Fisher), and subsequent analyzed using confocal microscopy (Zeiss LSM780).

