## Supplementary data for "Selective phosphorylation of AKT isoforms in response to dietary cues"

**Title:**

**Supplement:**

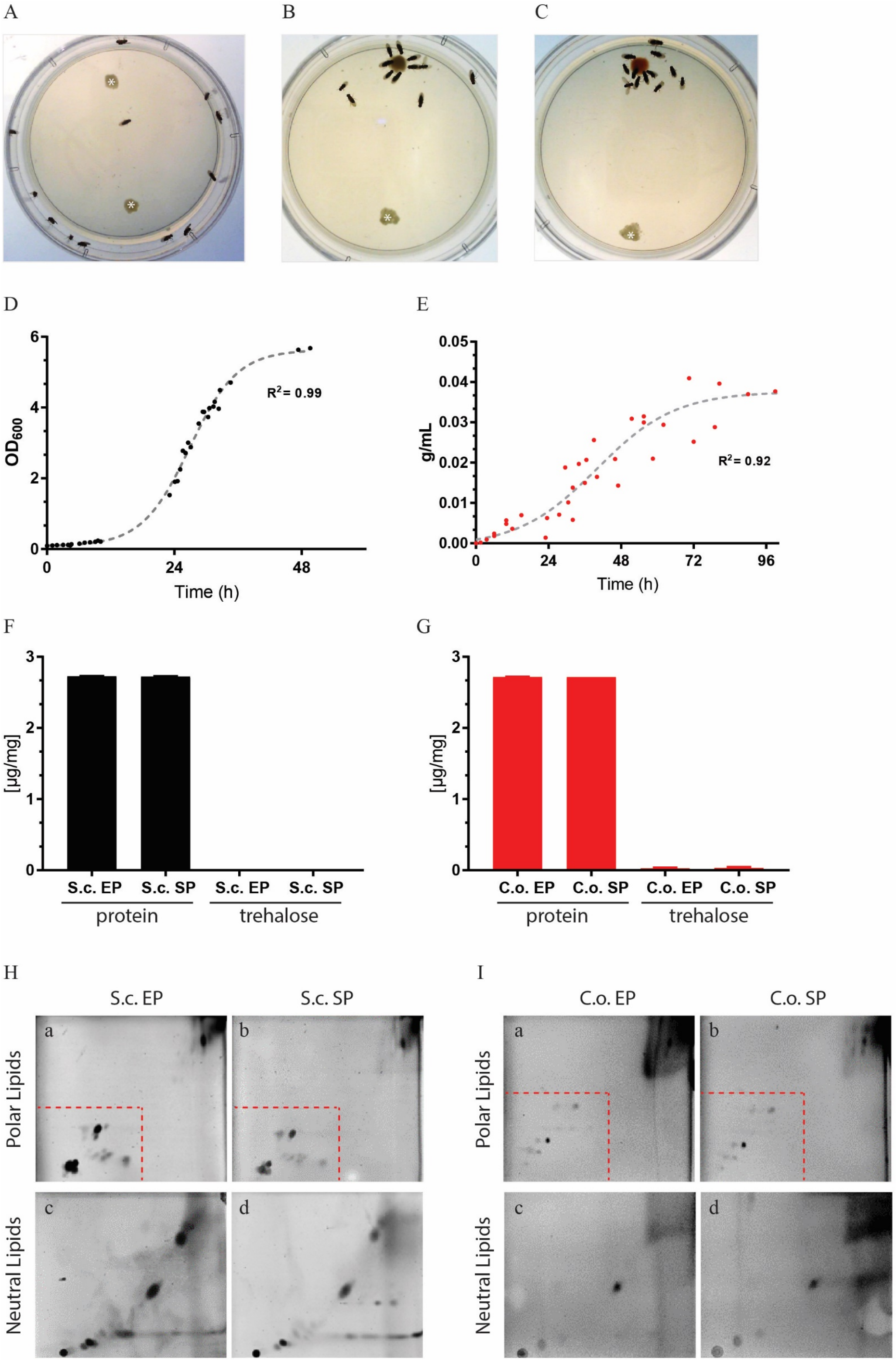


**Supplementary Figure 1: Yeast growth state is important for lipid quality in *S.cerevisiae***

A-C Shown are screenshots from movies testing fly's yeast attraction. Control plate probing for opposing plant food (*) baits (A), and assay plates testing *S.cerevisiae* (B) or *C.oligophagum* (C) against plant food bait.

D, E Plotted are the growth curves of *S.cerevisiae* (D) or *C.oligophagum* (E). $R^2$ indicates goodness of fit to non-linear sigmoidal regression model.

F, G Plotted are the protein and trehalose content of samples based on exponential (EP) and stationary (SP) *S.cerevisiae* (S.c.) in F or *C.oligophagum* (C.o.) in G.

H, I Lipid profiles of polar (Ha,b and Ia,b) and neutral lipids (Hc,d and Ic,d) based on exponential (EP) (Ha,c) and stationary (SP) (Hb,d) *S.cerevisiae* (S.c.) or *C.oligophagum* (C.o. EP (Ia,c), C.o. SP (Ib,d)) were analysed by 2-D Thin-layer chromatography. Red-dashed line isolated region is marked for Fig.2C,D.

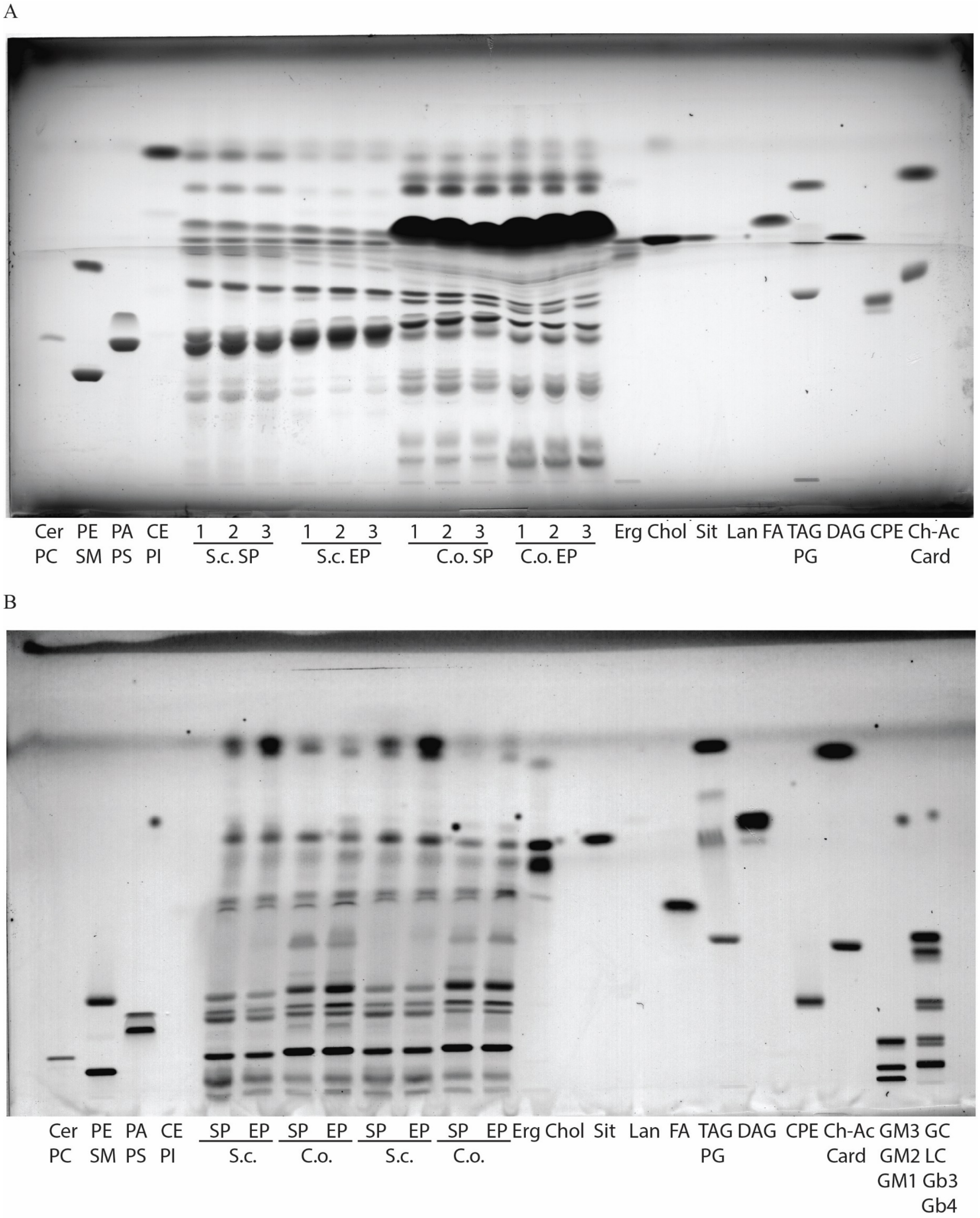


**Supplementary Figure 2: Heat-treatment changes the lipid profile**

Shown is the lipid profile of yeast and yeast food lipid extract samples by separation with 1-D Thin-layer chromatography. Samples are based on exponential (EP) and stationary (SP) *S.cerevisiae* (S.c.) or *C.oligophagum* (C.o.) in A, and respective foods in B. Lipid markers include: Card=cardiolipin, CE=cholesterol esters, Cer=ceramide, Ch-Ac=cholesterol-acetate, Chol=cholesterol, CPE=ceramide phosphorylethanolamine, DAG=diacylglycerol, Erg=ergosterol, FA=fatty acid, Gb3,4=globotriaosylceramide, GC=glucosylceramide, GM1-3=ganglioside, Lan=lanosterol, LC=lactosylceramide, PA=phosphatidic acid, PC=phosphatidylcholine, PE=phosphatidylethanolamine, PG=phosphatidylglycerol, PI=phosphatidylinositol, PS=phosphatidylserine, Sit=sitosterol, SM=sphingomyelin, TAG=triacylglycerol.

A

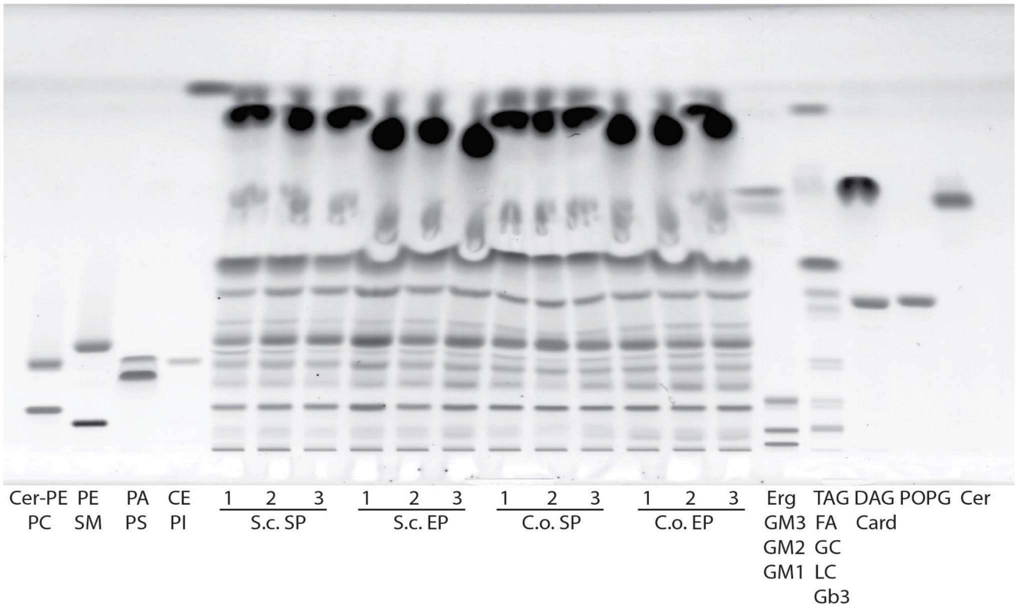


B

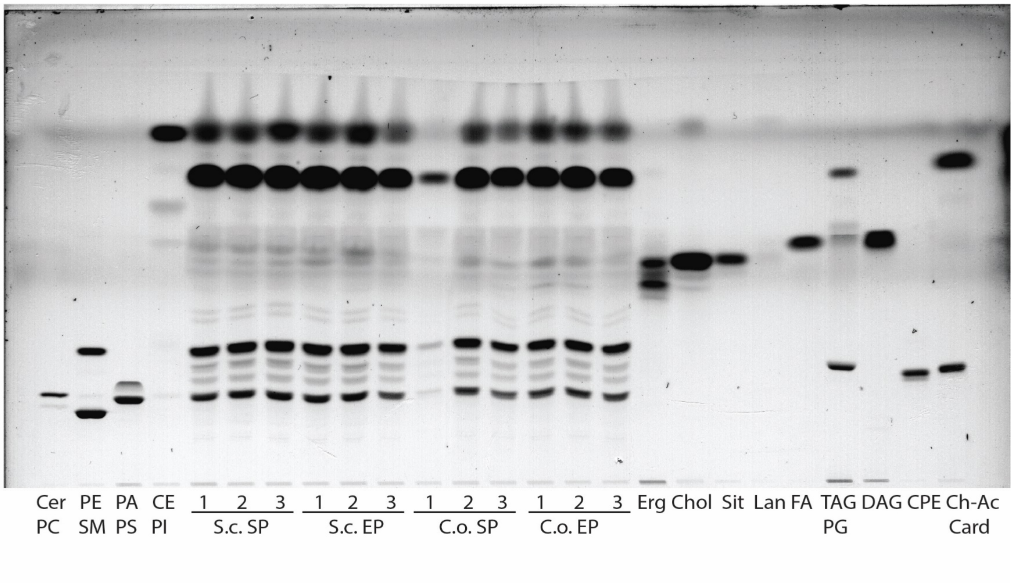


**Supplementary Figure 3: Fly endogenous lipid composition is not altered by nutritional lipids**
Shown is the separation of larval (A) and adult head (B) lipid extract samples by 1-D Thin-layer chromatography. Samples from animals kept on food based on exponential (EP) and stationary (SP) *S.cerevisiae* (S.c.) or *C.oligophagum* (C.o.). Samples were analysed in triplicates (1-3). Lipid markers include: Card=cardiolipin, CE=cholesterol esters, Cer=ceramide, Ch-Ac=cholesterol-acetate, Chol=cholesterol, CPE=ceramide phosphorylethanolamine, DAG=diacylglycerol, Erg=ergosterol, FA=fatty acid, Gb3=globotriaosylceramide, GC=glucosylceramide, GM1-3=ganglioside, Lan=lanosterol, LC=lactosylceramide, PA=phosphatidic acid, PC=phosphatidylcholine, PE=phosphatidylethanolamine, PG=phosphatidylglycerol, PI=phosphatidylinositol, POPG= 2-Oleoyl-1-palmitoyl-sn-glycero-3-phospho-rac-(1-glycerol), PS=phosphatidylserine, Sit=sitosterol, SM=sphingomyelin, TAG=triacylglycerol.
